# Analysis of Intermolt and Postmolt Transcriptomes Provides Insight into Molecular Mechanisms of the Red Swamp Crayfish, *Procambarus Clarkii* Molting

**DOI:** 10.1101/2020.12.09.418467

**Authors:** Shengyan Su, Brian Pelekelo Munganga, Can Tian, JianLin Li, Fan Yu, Hongxia Li, Meiyao Wang, Xinjin He, Yongkai Tang

**Author notes:** Contributed equally (Shengyan Su, Brian Pelekelo Munganga). Corresponding authors: Yongkai Tang.

## Abstract

In the present study, we explored expression changes in the transcriptomes of two molting stages (post-molt (M) and Intermolt (NM)) of the red swamp crayfish. A total of 307608398 clean reads, with an average length of 145bp were obtained. Further clustering and assembly of the transcripts generated 139100 unigenes. The results were searched against the NCBI, NR, KEGG, Swissprot, and KOG databases, in order to annotate gene descriptions, associate them with gene ontology terms, and assign them to pathways. Several genes and other factors involved in a number of molecular events critical for molting, such as energy requirements, hormonal regulation, immune response, and exoskeleton formation were identified, evaluated, and characterized. The information presented here provides a basic understanding of the molecular mechanisms underlying the crayfish molting process, with respect to energy requirements, hormonal regulation, immune response, and skeletal related activities during post-molt stage and the intermolt stage.

## 1. Introduction

Crustacean rearing has become an important sector of aquaculture (Bondad-Reantaso et al., 2012; Hosamani et al., 2017). Crustaceans are rich in protein and are an excellent source of minerals and vitamins (zinc, iron and Vitamin B-12, and choline, among others) compared to finfish (Venugopal, & Gopakumar, 2017; Hosamani et al., 2017). Given the rapid rate of human population growth around the world, crustacean’s contribution to meeting human protein needs will continue rising. However, the crustacean industry has its bottlenecks, which are disease outbreaks, cannibalistic behavior in addition to the lack of ways to enhance their growth (Allen & Steeby; Stentiford, 2012). Crustaceans undergo gradual growth which regularly requires the shedding of their exoskeleton, known as molting, to grow in size (Skinner, 1985). In many cases, molting is also necessary for copulation and successful reproduction in other crustaceans (Smith, & Ritar, 2008). This characteristic has significant implications to cannibalistic timing because after molting, the animals are unable to defend themselves and are hence highly vulnerable to cannibalism until their new shell is fully calcified. So far, the application of molecular techniques has solved several problems related to fisheries management, conservation, and aquaculture (Wenne et al., 2007; Mcandrew & Napier, 2011), therefore, studying the molecular molting mechanisms is imperative.

Red swamp crayfish (*Procambarus clarkii*) is an important kind of crustacean, and molting is a critical process in red swamp crayfish. Like other crustaceans, the red swamp crayfish molting process is divided into four hormone-controlled continuous phases; the intermolt, pre-molt, molt known as ecdysis, and post-molt (Skinner, 1985; Aiken & Waddy, 1987; Chang, 1995). Molting is a multifaceted process controlled by elaborate regulatory factors, including neuropeptide hormones and ecdysteroids (Naya, et at., 1989; Lachaise, et al, 1993; Pamuru, et al., 2012). These hormones are secreted by two endocrine glands, the paired Y-organs (glands in the maxillary somites) and neurosecretory cells (medulla terminalis X-organ sinus gland (XO-SG) complex) located within the eyestalks (Fingerman,1987; Taketomi, et al., 1993). The neurosecretory cell secretes neuropeptide hormone, a molt-inhibiting hormone (MIH) which prevents molting during the intermolt and postmolt stage (Skinner, 1985; Shyamal, et al., 2014). While the Y-organs, secrete ecdysteroids hormone (a derivative of ecdysone, 25-deoxyecdysone, and 3-dehydroecdysone) that stimulate molting. MIH circulation inhibits the synthesis of ecdysteroid by the Y-organs for most of the molt cycle (intermolt). Under favorable external and/or internal conditions (e.g temperature, light, loss of limbs) negative feedback is sent to the XO-SG that results in decreased circulation of MIH. Reduced MIH level stimulates synthesis and release of ecdysteroid by the Y-organ thereby initiating the premolt stage (Skinner, 1985; Keller, 1992). MIHs are not only involved in inhibiting synthesis and release of ecdysteroids but they have also been observed to play a role in reproduction (Jiang, et al., 2009; Wang, et al., 2014).

Earlier studies have led to a deeper understanding of the physiological and endocrine processes that take place during molting (Kleinholz, 1942; Fingerman,1987; Skinner, 1985; Lee, et al., 2017), thereby giving a chance for manipulation and control of the molting processes. So far molting in crustaceans can be induced by hormones or by eye-stalk ablation. Eye-stalk ablation shuts down all the hormones that inhibit the animal to undergo ecdysis (Venkitaraman et al., 2010; Lee, et al., 2017; Rana, 2018). While hormonal control (melatonin, α-ecdysone e.t.c.), suppresses the production of MIH, which in turn initiates the production of ecdysteroids thereby leading to molting (Diwan, 2020). Although the physiological and endocrinal mechanisms of molting have been widely studied, a lot regarding molecular mechanisms of molting in crustaceans is yet to be revealed and learned. Moreover, studies on crayfish molecular molting mechanisms are still few, hence, these mechanisms still remain poorly understood.

To the best knowledge of the authors, only the expression of cytoskeletal and molt-related genes has been conducted to elucidate the molecular mechanisms underlying the red swamp crayfish molting process (Tom, et al., 2014). In that study, Tom and colleagues (2014), identified a number of genes and other factors related to exoskeleton formation and the major transcriptional events during premolt and their timing was determined. Nevertheless, molecular molting mechanisms have been studied in other crustaceans and several molting related factors have been characterized (Kuballa, et al., 2011; Gao et al., 2015; Huang, et al., 2015; Shyamal, et al., 2018). For example, Gao et al. (2015), identified several genes related to immunity, hormonal regulation, and other factors related to molting when they conducted a whole transcriptome analysis of the molting process. Kuballa et al. (2011), identified factors related to energy cellular energy requirements cycle, cuticular protein, hemocyanin, cuticle hardening, muscle formation, and lipid metabolism across the different stages of the molting process.

In order to understand the mechanisms and molecular events related to crayfish molting, RNA-Sequencing (RNA-seq) was used to explore the expression changes of genes that occur between the intermolt (NM) stage and postmolt (M). Tissues from the hepatopancreas of the crayfish in the two molting stages were used for transcriptome sequencing. This work will contribute to a better understanding of the crustacean molting regulatory mechanisms thereby shedding light on how molecular mechanisms can be used to intervene in the molting process for improved culture and management of red swamp crayfish and other crustaceans. Furthermore, our work provides a valuable resource of crayfish transcriptome genome annotation for further identification of candidate genes controlling molting in crayfish and other molting animals.

## 2. Materials and Methods

### 2.1 Experimental Animals

A total of 40 red swamp crayfish weighing 10-25g were obtained from the greenhouse to the experimental laboratory at Fresh Water Research Center of the Chinese Academy of Fishery Science (FFRC). The red swamp crayfish were put in 4 equal size glass tanks at a density of 10 crayfish per tank and acclimated to the laboratory conditions for two weeks. After two weeks the molting process was observed 24hrs daily and the crayfish were fed with commercial feed to satiation (three times a day; 9 AM, 3 PM, and 7 PM). The crayfish were cared for in accordance with the international and China’s guide for the care and use of experimental animals.

### 2.2 Sample Collection and Preparation

15 minutes after the molting process was observed (post molt stage, designated as M), the red swamp crayfish were sampled and dissected to collect the hepatopancreas, at the same time the crayfish in the intermolt (non-molting NM)) was also dissected to collect the hepatopancreas. The collected tissue from both crayfish in the two molt stages (post molt and intermolt stage) were fixed and stored in a −80 degrees freezer awaiting RNA extraction. A total of 6 crayfish were sampled, 3 in the post molt stage (M) and the other 3 in the intermolt stage (NM).

### 2.3 RNA isolation and RNA-Seq library preparation

Approximately 80mg of hepatopancreas tissue was used for RNA extracted using Trizol (Invitrogen, Carlsbad, CA, USA) following the manufacturer’s instructions. The quality and amount of RNA integrity were verified using an Agilent 2100 Bioanalyzer (Agilent, Shanghai, China). Following the manufacturer’s instruction 3μg RNA per group was used for sequencing libraries were generated using a NEBNext Ultra RNA Library Prep Kit for Illumina (NEB, USA). The index-coded samples were clustered on a cBot Cluster Generation System using the TruSeq PE Cluster Kit v3-cBot-HS (Illumina) as outlined by the manufacturer. Then, on an Illumina Hiseq 2000 platform, the library preparations were sequenced and 100 bp single-end reads were obtained.

### 2.3 Transcriptome assembly and annotation

The raw reads obtained after sequence were screened for adaptor-polluted, low-quality, high content of unknown base (N) reads, non-coding RNA (such as rRNA, tRNA and miRNA) and empty reads, to obtain clean reads. The transcripts with length of at least 100 bp were used for further analysis. The clean reads were assembled using trinity (v2.0.6) (Vasani, N. (2019). into 272346 transcripts with GC content of was 47.71% and 46.80%. The transcripts length ranged from 200 to over 3000 bp, with an average of of 885bp and an N50 of length of 1806bp (Table 2).

Using BLASTX against NCBI, NR, KEGG, Swissprot, and KOG databases, assembled transcriptomes were annotated, with a cutoff E-value lower than 1e-6. BLASTX findings were imported into the BLAST2GO program, then the EC numbers for the KEGG pathway and Gene Ontology (GO) terms were annotated (Conesa, A. et al).

### 2.4 GO and KEGG analysis of differentially expressed Genes

ClusterProfiler (Yu et al., 2012), was used to classify the overrepresented GO terms in the biological process (BP), molecular function (MF) groups and cellular component (CC), as well as the KEGG pathway categories, to determine the functions and significantly enriched pathways of the DEGs. For these analyses, the hypergeometric distribution threshold was a p-value of < 0.05.

### 2.5 qRT-PCR

To validate RNA-seq data and expression, we chose eight genes that were significant enriched in KEGGY pathway analysis and **AF436001** was used as an internal reference gene. The primers for the selected genes were designed using designed using Primer3 Input (v.0.4.0), (https://primer3.ut.ee/) (Tab. S1). Approximately, 2 μg of total RNA per sample was used to generate first-strand cDNA using reverse transcription system (Takara, Dalian, China). Quantitative RT-PCR was performed using SYBR Premix Ex Taq (Takara, Dalian, China) on ABI 7500 system. The amplification program was performed as 95 °C for 2 min, followed by 95°C for 15 s and 60°C for 31 s (40 cycles). Three biological replicates were performed for each gene. The C). The relative expression levels of genes were calculated using the 2^-CT^ method (Livak & Schmittgen, 2001).

## 3. Results

### 3.1 Ecdysteroid Changes in Crayfish Morphology

Figure 1 shows the ecdysteroid changes that occur in the morphology of crayfish. In our study we observed the molting process as it proceeded from the intermolt stage through to the postmolt stage. In the intermolt stage the crayfish had a fully developed hard exoskeleton (Fig. 1**a**), then the exoskeleton was partially digested during the premolt staging as it progressed towards molting. In the molt stage water was absorbed to increase the body volume leading to the splitting of the partially degraded old exoskeleton (Fig. 1**c**). After molting the crayfish was very soft and vulnerable due to the exoskeleton which was not calcified, it is at this stage that the exoskeleton and membranous layers begin to be formed (Fig. 1**b**).

**Figure 1.**
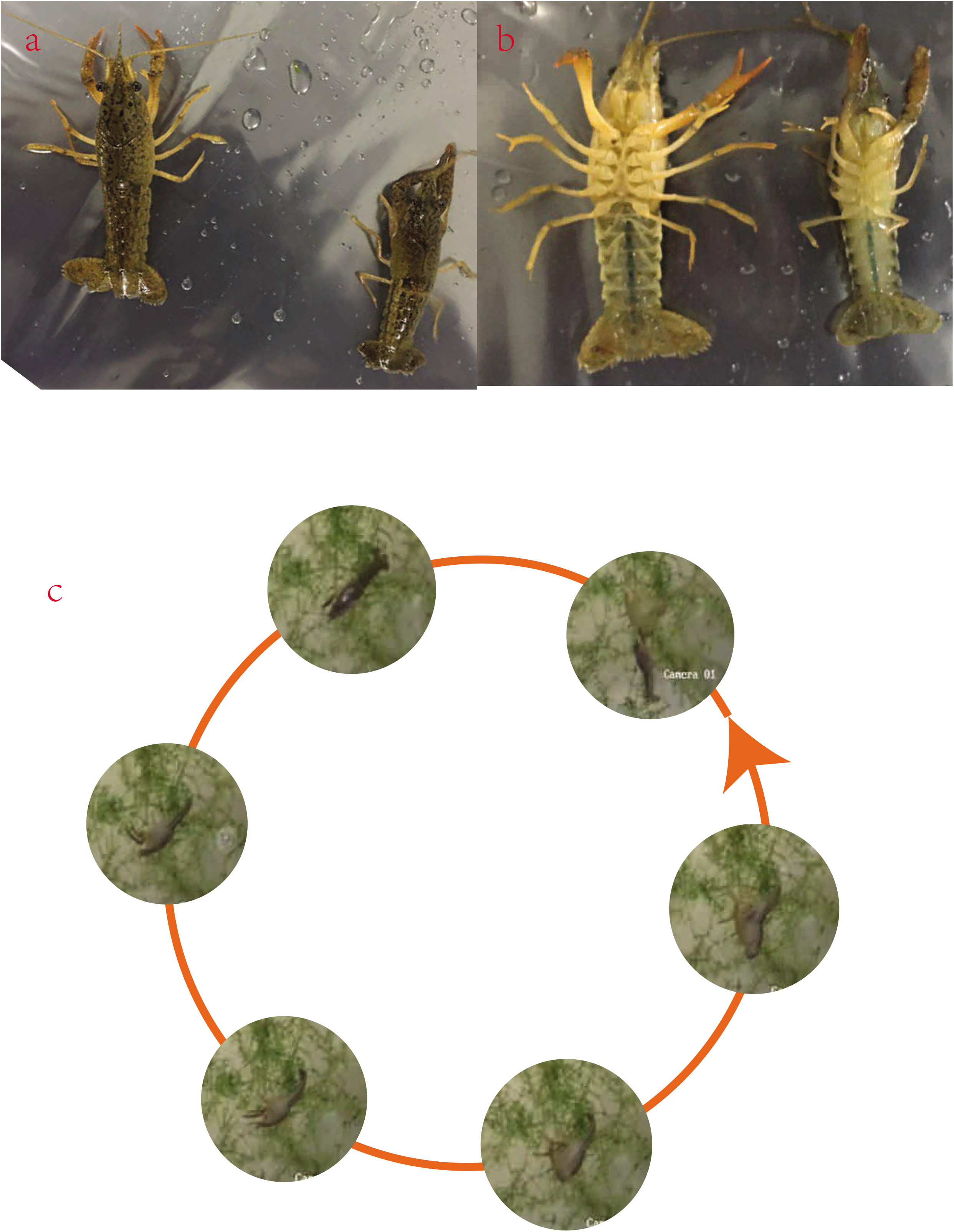
Shows the morphological changes that occur during the molting process: (a) Inter-molt stage (b) Post-molt stage (c) sequence of the molting process.

### 3.2 Read Sequencing, Assembly, and Mapping

For this study the transcriptomes we obtained from the hepatopancreas of the post-molt (M) and the non-molt (NM) crayfish. The acquired raw reads were trimmed and sifted using Illumina sequencing platform, generating a total of 307608398 clean reads, with an average length of 145bp (Table 1). The clean reads were deposited into the Sequence Read Archive database at NCBI (PRJNA592222). The average GC content of the clean reads was 47.71%, while the GC% for NM (48.61%) was slightly higher than that of the M (46.80%). The proportion of nucleotides with quality values larger than 30 in reads (Q30) was 95.06%, with the NM representing 94.60% and M representing 95.51% (Table 1).

Using Trinity program, a total of 272346 transcripts were assembled from the clean reads, having an average length of 885bp and an N50 of length of 1806bp. Of these transcripts 41.5% were more than 500bp in length and 24.3% longer than 1000bp (Table 2). Further clustering and assembly of the transcripts generated 139100 unigenes. These unigenes (139100) had an average length of 676bp, of which 33.1% were longer than 500bp in length and 24.3% longer than 1000bp (Table 2, Fig. S1**b**). Furthermore, most of the identified DEGs had length between 200-300, GC content within the range 30-50%, and Isoforme ranging between 2-3 (11.4%) (Fig. S1). To assess the extent of coverage of the assembled unigenes and evaluate the effects of coverage depth on unigene assembly, the percentile of gene body against coverage depth was calculated and plotted (Fig. S2**a**).

The clean reads for the two libraries (M and NM) were aligned against the reference genome database, in which we found that 94.7% of the M reads and 95.7% of the NM reads were mapped to the reference genome. Among the M reads (94.7%), 76.0 % were multi-mapped, 18.7% uniquely mapped, and 16.6% were mapped in proper pairs. Of the NM reads (95.7%) mapped, 77.3 % were multi-mapped, 18.4% uniquely mapped, and 15.9% were mapped in proper pairs. Splice reads for both libraries was 0%, while non-splice reads were 18.7% and 18.4% for M and NM respectively (Tab. 3). The assessment of unique gene mapping ratio and the genome mapping ratio met the requirements for follow-up analysis.

### 3.3 Functional Annotation

The Assembled unigene were searched against the NCBI, NR, KEGG, Swissprot, and KOG databases, using the BLASTx (E-value, 10^25^). The annotation results showed that out of the 139100 assembled unigenes, 5064 unigenes had homologous sequences in all the four databases; 96 unigenes in the KEGG, Swissprot, and KOG; 625 in the NR, KEGG, and Swissprot; and 3961 in the NR, Swissprot, and KOG. Furthermore, we found out that 6511 unigenes only had homologous sequences in NR, 5 unigenes only in KEGG, 570 unigenes only in Swissprot, and 74 unigenes only in KOG. The total number of unigenes with homologous sequences in each database were, 19217 in NR, 5863 KEGG, 13484 Swissprot, and 9509 in the KOG database. The annotation results are illustrated in the Venn diagram (Tab. S2 & Fig S2**b**).

### 3.4 Characterization of SSR, SNPs, and InDels

The simple sequence repeats (SSR) ranged from one to six. The number of SSRs with each repeat unit was found to vary, the most common SSRs were mono-nucleotide repeats, followed by di-nucleotide repeats and the hexa-nucleotide repeats were the least common (Fig. S3**a**). The SNPs and InDels distributions within various genomic features were revealed by annotation of the transcripts from all samples to the crayfish reference genome. Generally, a similar distribution pattern in both SNPs and InDels was observed in the six samples (Fig. S3**b**). The SNPs and InDels were highest in M2, followed by M1, and were least in NM1. The average number of the M SNPs were higher than that of NM.

### 3.5 Differentially Expressed Genes Analysis

DEGs analysis of the two libraries, showed that 130584 DEGs were expressed in the M stage (94.6% of the genes) and 116754 were expressed in the NM stage (84.6% of the genes), among which 109321 were expressed in both the M and NM stages. Furthermore, 21263 were only expressed in the M, while 7433 were only expressed in NM (Fig. 2**a**). To understand and illustrate the relationship between the M and NM stages, we performed a hierarchical clustering of the DEGs from all the samples, using Ward’s method of Euclidean distances (Ward, 1963) (Fig. 2**b**). The results showed a higher correlation between samples in each molting stage (M and NM), hence, big differences occur between the two molting stages. Similarly, *Principal Component* Analysis (PCA), plot revealed that the DEGs from the all samples could be clustered into two groups (M and NM), based on similarities of gene expression patterns (Fig. 2**c**). In addition, a dendrogram placed the sample DEGs into two main clusters; M1, M2 and M3 into one cluster and NM1, NM2, and NM3 into another cluster (Fig. 2**d**). Similar trends, were observed in the box plot of the sample DEGs (Fig. 2.**e** & **f**). A heatmap was plotted to illustrate the relative expression profiles of the DEs in the two molting stages (M and NM) (Fig. 3**a**). There were two general profiles observed, many of DEs showed highest relative expressions in N stage (profile 2), compared to the NM stage (profile 1). Among the identified DEGs, 1401 were up regulated and 946 were down regulated (Tab **S3** & Fig. **3b**). The volcano plots present the distribution of the differentially expressed genes, according to the FDR ≤ 0.001 and Log Fold change threshold (3**c**).

**Figure 2.**
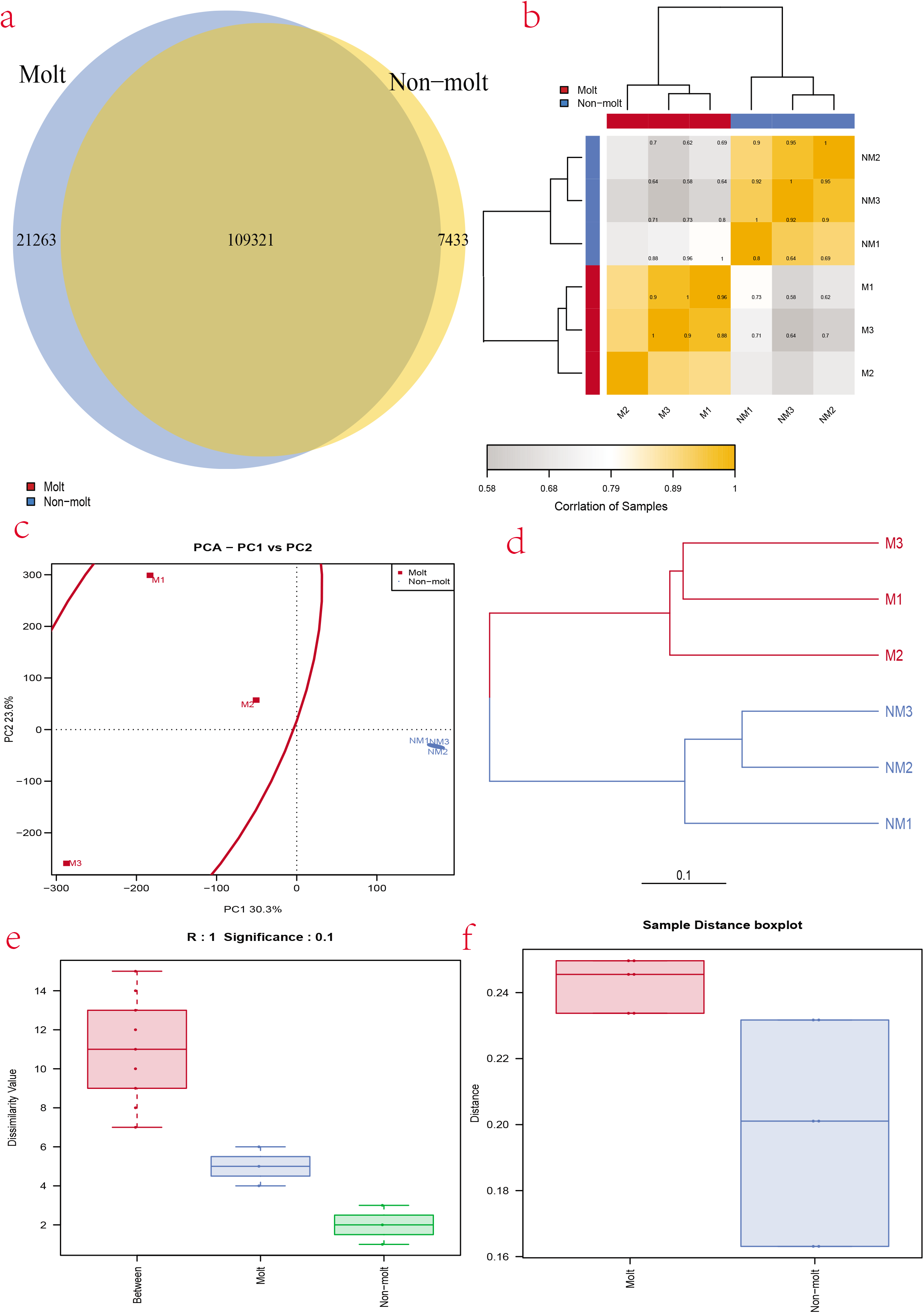
(a) Venn diagram showing the number of DEGs expressed in each molting stage (N&NM) and those expressed in both stages. (b) The DEGs correlation heatmap of sample clustering. The more similar the two samples are, the nearer the distance is. Each color box represents the distance between different samples. The greater the distance, the whiter and the closer the distance, the deeper the yellow. (c) Cluster analysis of the samples: PCA plot is illustrating the similarities and difference of the samples from the two molting stags (M and NM). (d) Cluster analysis of the samples: dendrogram is the is illustrating similarities and difference of the samples from the two molting stags. (e) Dissimilarity box plot illustrating the relationship between samples from the two molting stages. (f) Distance box plot illustrating the relationship between samples from the two molting stages.

**Figure 3.**
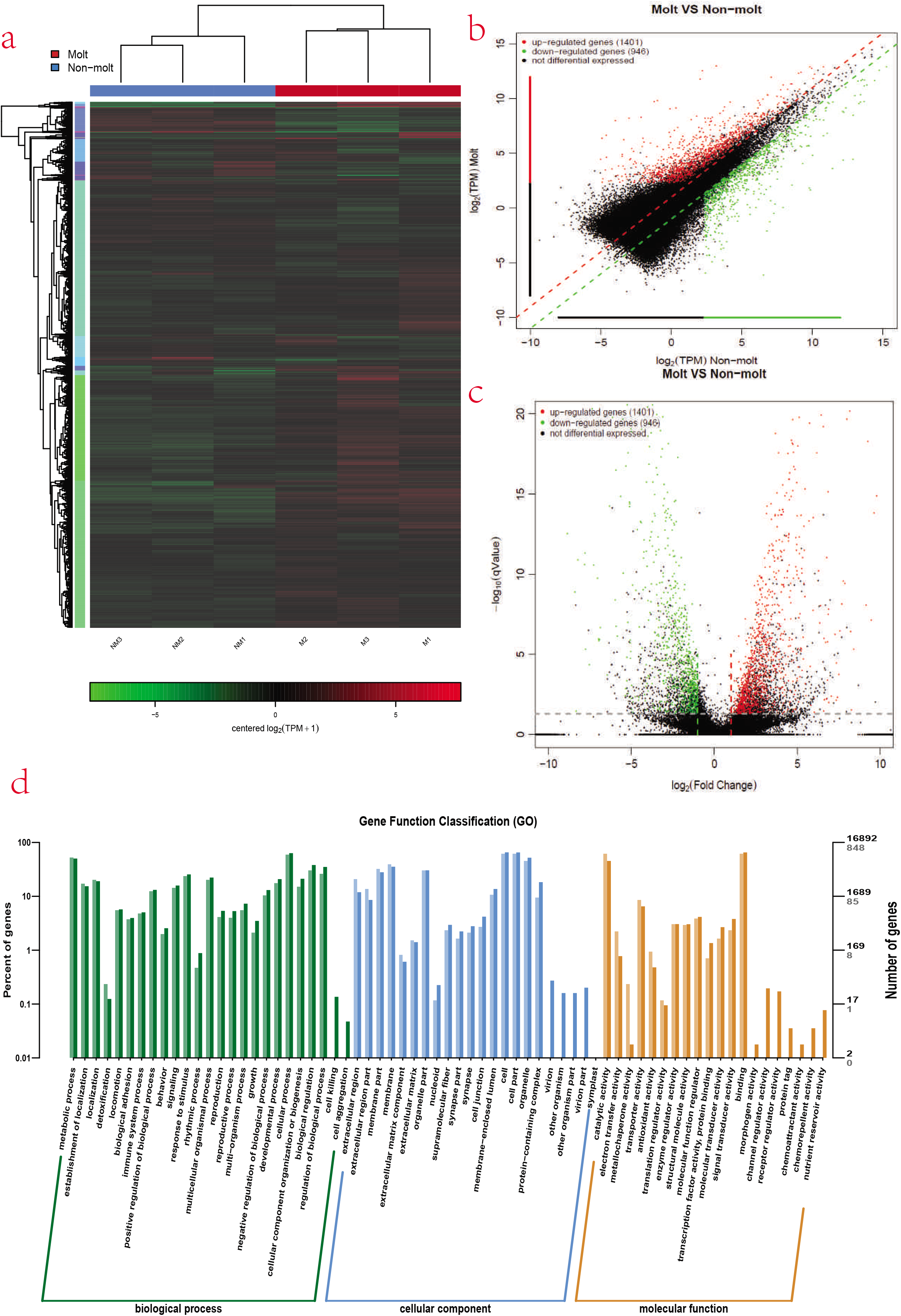
**(a)** Heat map of differentially expressed genes, Colored keys represent the fold changes (log2 transformed counts) of gene expression between the molt stages; Red represents up-regulation and green represents down-regulation. Each column represents moltign stage. (b) Volcano plot showing log2 (TPM) Non-molt (x-axis) and log2 (TPM) Molt (y-axis) of the differentially expressed genes, where red represent up-regulation, green downregulation and black represent non-differentiated genes. (c) Volcano plot showing log2 (fold change) (x-axis) and significance (−log_10_ * adjusted p-value; y-axis) of the differentially expressed genes, where red represent up-regulation, green downregulation and black represent non-differentiated genes. (d) Gene ontology (GO) classification of transcripts from the two samples (N and NM). The three main GO categories include biological process (blue), cellular component (red), and molecular function (green).

### 3.6 GO and KEGG analysis of differentially expressed genes

To acquire complete functional information and classification of M vs NM, unigenes were aligned against GO and KEGG databases. Gene Ontology (GO) analysis, revealed that 16892 unigenes were annotated. These unigenes were sorted into 67 functional groups which belong to GO categories including biological process, cellular process, and molecular function. In the cellular component category, the largest cluster of DEGs were associated with cell and cell part. Under the biological process category, abundant DEGs were involved in cellular process, development process, and biological process. Within the molecular function category, a large proportion were associated with catalytic activities and transporter activity (Fig. 3**d**). A heat map which grouped genes according to FPKM values was generated in Cluster (de Hoon et al. 2004) and visualized in TreeView to analyse their expression levels across molting stages (Saldanha 2004), (Fig. 3a).

To further understand the biological pathways of the genes it is necessary to carry out a pathway-based analysis (KEGG), (Cirillo, et al., 2017). For this purpose, the 5863 unigenes were mapped to the reference of typical pathways in the KEGG database (Fig. 4**b**). (KEGG), analysis, revealed that 5863 unigenes were annotated. The DEGs were mapped to 33 different pathways, belonging to five KEGG pathways: cellular process, environmental processing information, genetic processing information, metabolism, and organismal system. To further understand the biological processes related to the identified pathways of various DEGs, KEGG enrichment analysis was conducted. A total of 30 KEGG processes were significantly enriched. The top ten KEGG pathways of DEGs were enriched in Starch and sucrose metabolism (ko00500), Lysosome (ko04142), Peroxisome (ko04146), Retinol metabolism (ko00830), Amino sugar and nucleotide ko00520), *Pentose and glucuronate* interconversions (ko00040), Longevity regulating pathway (ko04212), Metabolism of xenobiotics by cytochrome P45 (ko00980, Fatty acid degradation (ko00071), and Tyrosine metabolism ko00350) (figure 4**a**). Moreover, understanding cellular function needs knowledge of functional interactions among the expressed proteins. Thus, we carried out STRING protein interaction network analysis, thereby producing networks from the up-regulated genes of M vs NM (figure 4**c** and **d**). Similar to KEGG enrichment, significantly enriched biological processes could be identified on the networks by degree score (−log_10_ (P value)).

**Figure 4.**
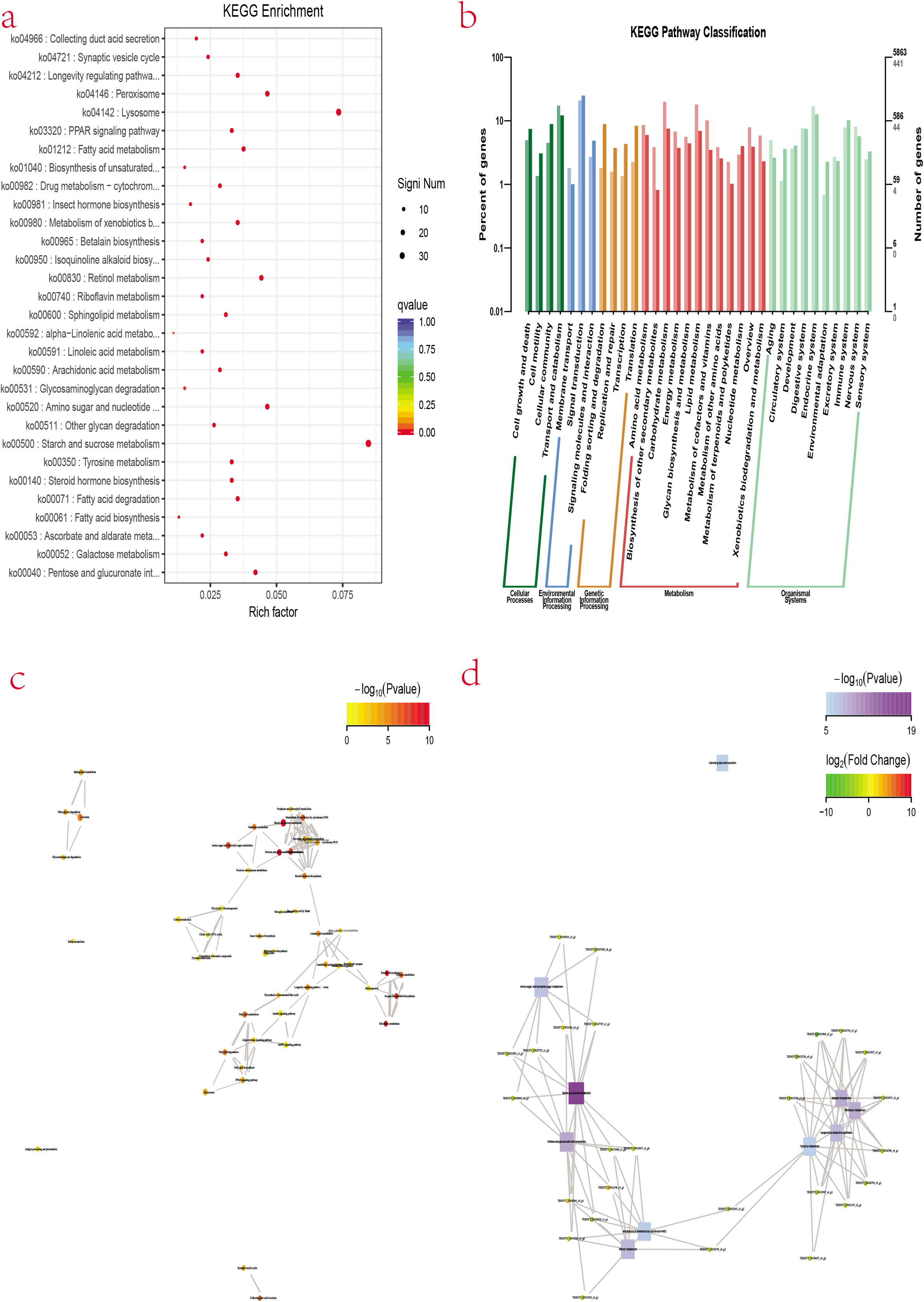
**(a)** Figure 4 Scatter plot of enriched KEGG pathways statistics. Rich factor is the ratio of the differentially expressed gene number to the total gene number in a certain pathway. Q-value is corrected P-value ranging from 0 ~ 1. The color and size of the dots represent the range of the Q-value and the number of DEGs mapped to the indicated pathways, respectively. Top 30 enriched pathways are shown in the figure. (b) Plots showing categories of genes classified based on the Kyoto Encyclopedia of Genes and Genomes (KEGG) analysis, five categories were identified. STRING interaction network analysis. (c) Proteins and metabolic pathway interactions. (d) Gene interaction network.

To learn about the dynamics of energy during the postmolt (M) and intermolt stage of the molting, factors related to metabolism were identified. Five, energy related factors including NADH dehydrogenase, Cytochrome p450, thyroid hormone signaling pathway, creatine kinase, arginine kinase were identified. The expression of energy related factors are detailed in Table 4. Among the identified energy related factors only Cytochrome and Thyroid hormone signaling pathway were down regulated in the M stage. The substantial variation in the expression of these factors in the two molting stages could be related with the fluctuating energy demands across the molt cycle.

**Tab. 4.** Factors Related to Energy Requirements During Molting

| Factor Name | Number of Genes | Gene ID | log2Fold Change | Expression Pattern |
| --- | --- | --- | --- | --- |
| NADH dehydrogenase | 3 | TRINITY_DN33948_c1_g8 | 1.768285 | Up-Regulated |
|  |  | TRINITY_DN29179_c0_g1 | 1.451322 | Up-Regulated |
|  |  | TRINITY_DN33884_c4_g8 | 1.945742 | Up-Regulated |
|  |  | TRINITY_DN33884_c4_g4 | 1.637054 | Up-Regulated |
| Cytochrome P450 | 8 | TRINITY_DN29218_c1_g1 | -1.49657 | Down-Regulated |
|  |  | TRINITY_DN29218_c1_g2 | -1.37342 | Down-Regulated |
|  |  | TRINITY_DN29218_c1_g3 | -1.27608 | Down-Regulated |
|  |  | TRINITY_DN29218_c1_g5 | -1.11205 | Down-Regulated |
|  |  | TRINITY_DN28462_c2_g2 | -1.56713 | Down-Regulated |
|  |  | TRINITY_DN33678_c0_g1 | -2.41627 | Down-Regulated |
|  |  | TRINITY_DN34018_c2_g1 | -1.26278 | Down-Regulated |
|  |  | TRINITY_DN33391_c6_g1 | 1.416798 | Up-Regulated |
| Thyroid hormone signaling pathway | 16 | TRINITY_DN32933_c1_g3 | -1.42248 | Down-Regulated |
|  |  | TRINITY_DN27599_c3_g1 | -2.45796 | Down-Regulated |
|  |  | TRINITY_DN27599_c5_g4 | -1.4136 | Down-Regulated |
|  |  | TRINITY_DN32047_c0_g1 | 1.822958 | Up-Regulated |
|  |  | TRINITY_DN29637_c1_g1 | 1.450007 | Up-Regulated |
|  |  | TRINITY_DN32340_c4_g2 | 1.571665 | Up-Regulated |
|  |  | TRINITY_DN32656_c5_g2 | -1.06408 | Down-Regulated |
|  |  | TRINITY_DN25356_c5_g2 | 2.010127 | Up-Regulated |
|  |  | TRINITY_DN32500_c3_g3 | 1.711267 | Up-Regulated |
|  |  | TRINITY_DN31858_c11_g1 | 1.577703 | Up-Regulated |
|  |  | TRINITY_DN32933_c1_g3 | -1.42248 | Down-Regulated |
|  |  | TRINITY_DN27599_c3_g1 | -2.45796 | Down-Regulated |
|  |  | TRINITY_DN27599_c5_g4 | -1.4136 | Down-Regulated |
|  |  | TRINITY_DN32047_c0_g1 | 1.822958 | Up-Regulated |
|  |  | TRINITY_DN29637_c1_g1 | 1.450007 | Up-Regulated |
|  |  | TRINITY_DN28273_c1_g2 | 1.929502 | Up-Regulated |
| Creatine kinase | 1 | TRINITY_DN25935_c2_g2 | 1.555388 | Up-Regulated |
| Arginine kinase | 1 | TRINITY_DN25935_c2_g2 | 1.555388 | Up-Regulated |
|  |  | TRINITY_DN27099_c7_g1 | -3.718 | Down-Regulated |

To reveal the expression fluctuations of the immune-related factors between the two molting stages (M and NM), we searched and evaluated molt-related immune factors identified and characterized in other arthropods with more focus on crustaceans. Approximately, nine immune-related factors including innate immune response, lysosome, toll-like receptors, caspase, trypsin-like serine protease, peroxiredoxin, thioredoxin, glutaredoxin and c-type lectin-like were identified (Tab. 5). Interestingly, majority of these immune-related factors were up-regulated.

**Tab. 5.** Immune Related Factors

**Tab 5.** Immune Related Factors
| **Factor Name** | **Number of Genes** | **Gene ID** | **log2Fold Change** | **Expression Pattern** |
| --- | --- | --- | --- | --- |
| **Innate immune response ()** | 3 | TRINITY_DN30285_c3_g1 | 2.426348 | Up-Regulated |
|  |  | TRINITY_DN31835_c3_g1 | 1.645295 | Up-Regulated |
|  |  | TRINITY_DN27554_c0_g1 | 1.760783 | Up-Regulated |
| **Lysosome** | 1 | TRINITY_DN29507_c1_g1 | 2.764554 | Up-Regulated |
| **Toll-like receptors** | 4 | TRINITY_DN27554_c0_g1 | 1.760783 | Up-Regulated |
|  |  | TRINITY_DN32340_c4_g2 | 1.571665 | Up-Regulated |
|  |  | TRINITY_DN28208_c1_g2 | 1.583255 | Up-Regulated |
|  |  | TRINITY_DN30049_c3_g1 | 2.06495 | Up-Regulated |
| **Caspase** | 3 | TRINITY_DN29822_c0_g1 | 1.706101 | Up-Regulated |
|  |  | TRINITY_DN30712_c4_g1 | 2.496235 | Up-Regulated |
|  |  | TRINITY_DN30639_c3_g2 | 2.232428 | Up-Regulated |
| **Trypsin-like serine protease** | 3 | TRINITY_DN27214_c1_g3 | 1.396213 | Up-Regulated |
|  |  | TRINITY_DN30994_c2_g1 | -1.17476 | Down-regulated |
|  |  | TRINITY_DN26076_c3_g1 | 3.835115 | Up-Regulated |
| **Peroxiredoxin** | 1 | TRINITY_DN27705_c3_g1 | 1.778722 | Up-Regulated |
| **Thioredoxin** | 3 | TRINITY_DN27258_c3_g2 | -1.958 | Down-regulated |
|  |  | TRINITY_DN27705_c3_g1 | 1.778722 | Up-Regulated |
|  |  | TRINITY_DN28159_c5_g1 | 1.880269 | Up-Regulated |
| **Glutaredoxin** | 1 | TRINITY_DN27705_c3_g1 | 1.778722 | Up-Regulated |
| **C-type lectin-like** | 6 | TRINITY_DN31486_c0_g1 | 1.601822 | Up-Regulated |
|  |  | TRINITY_DN34044_c3_g1 | 2.00469 | Up-Regulated |
|  |  | TRINITY_DN32410_c3_g3 | 3.095355 | Up-Regulated |
|  |  | TRINITY_DN31344_c4_g2 | -3.12709 | Down-Regulated |
|  |  | TRINITY_DN25837_c0_g1 | 2.852943 | Up-Regulated |
|  |  | TRINITY_DN27099_c7_g1 | -3.718 | Down-Regulated |

Hormones are important regulators of red swamp crayfish molting cycle and they involved are in numerous process such as osmoregulation, reproduction, glucose metabolism e.t.e (Shyamal, et al., 2014; Wang, et al., 2014). To understand the global changes in hormone regulation system that occur as the molt process progresses, we enumerated several hormonal related factors. As a result, we identified nine types of molting factors related to hormone regulation (Tab. 6). Three of these factors were upregulated and the other three were down-regulated. Two genes related to MIH and crustacean hyperglycemic hormone (CHH) were identified.

**Tab. 6.** Hormonal Regulations Related Factors

**Tab 6. Hormonal Regulations Related Factors**
| **Factor Name** | **Number of Genes** | **Gene ID** | **log2Fold Change** | **Expression Pattern** |
| --- | --- | --- | --- | --- |
| **MIH** | 2 | TRINITY_DN28405_c2_g1 | 2.124327 | Up-Regulated |
|  |  | TRINITY_DN28405_c2_g3 | 2.093657 | Up-Regulated |
| **Crustacean hyperglycemic hormone (CHH)** | 2 | TRINITY_DN28405_c2_g1 | 2.124327 | Up-Regulated |
|  |  | TRINITY_DN28405_c2_g3 | 2.093657 | Up-Regulated |
| **Hyperglycemic hormone-like peptide 2 precursor** | 2 | TRINITY_DN28405_c2_g3 | 2.093657 | Up-Regulated |
|  |  | TRINITY_DN28405_c2_g1 | 2.124327 | Up-Regulated |
| **Neuroparsin** | 2 | TRINITY_DN30757_c1_g2 | -3.81307 | Down-regulated |
|  |  | TRINITY_DN9917_c0_g1 | 2.370687 | Up-Regulated |
| **Ecdysteroid regulated-like protein** | 1 | TRINITY_DN29775_c2_g1 | 2.405408 | Up-Regulated |
| **Ecdysteroid kinase** | 5 | TRINITY_DN33260_c1_g1 | -2.36332 | Down-regulated |
|  |  | TRINITY_DN27997_c3_g2 | -2.71021 | Down-regulated |
|  |  | TRINITY_DN33944_c2_g5 | -2.02487 | Down-regulated |
|  |  | TRINITY_DN32865_c4_g2 | 8.170282 | Up-Regulated |
|  |  | TRINITY_DN30786_c4_g3 | -2.6306 | Down-regulated |
| **Vitellogenin** | 1 | TRINITY_DN29089_c2_g1 | -4.02228 | Down-regulated |
| **17-beta-dehydrogenase 8-like isoform X1** | 3 | TRINITY_DN33761_c1_g1 | -2.73846 | Down-regulated |
|  |  | TRINITY_DN23109_c0_g1 | -1.32568 | Down-regulated |
|  |  | TRINITY_DN31520_c0_g1 | -3.07782 | Down-regulated |
| **E75 nuclear receptor** |  | TRINITY_DN27618_c2_g1 | 2.65079 | Up-Regulated |

Expression patterns of factors implicated in exoskeleton formation were investigated to identify those that could be possibly be involved in the molting process. By data mining of the annotated transcripts with reference to previous studies of exoskeleton formation related factors such as C-type lectin-like, manose-binding protein, *Eriocheir sinensis* chitin synthase gene, CDA like 2, CBM 14, and cryptocyanin 1 identified (Tab. 7). The expression profile of majority factors showed a characteristic pattern of up-regulation at M stage compared to NM. However, cryptocyanin 1was down regulated at m stage.

**Tab. 7.**
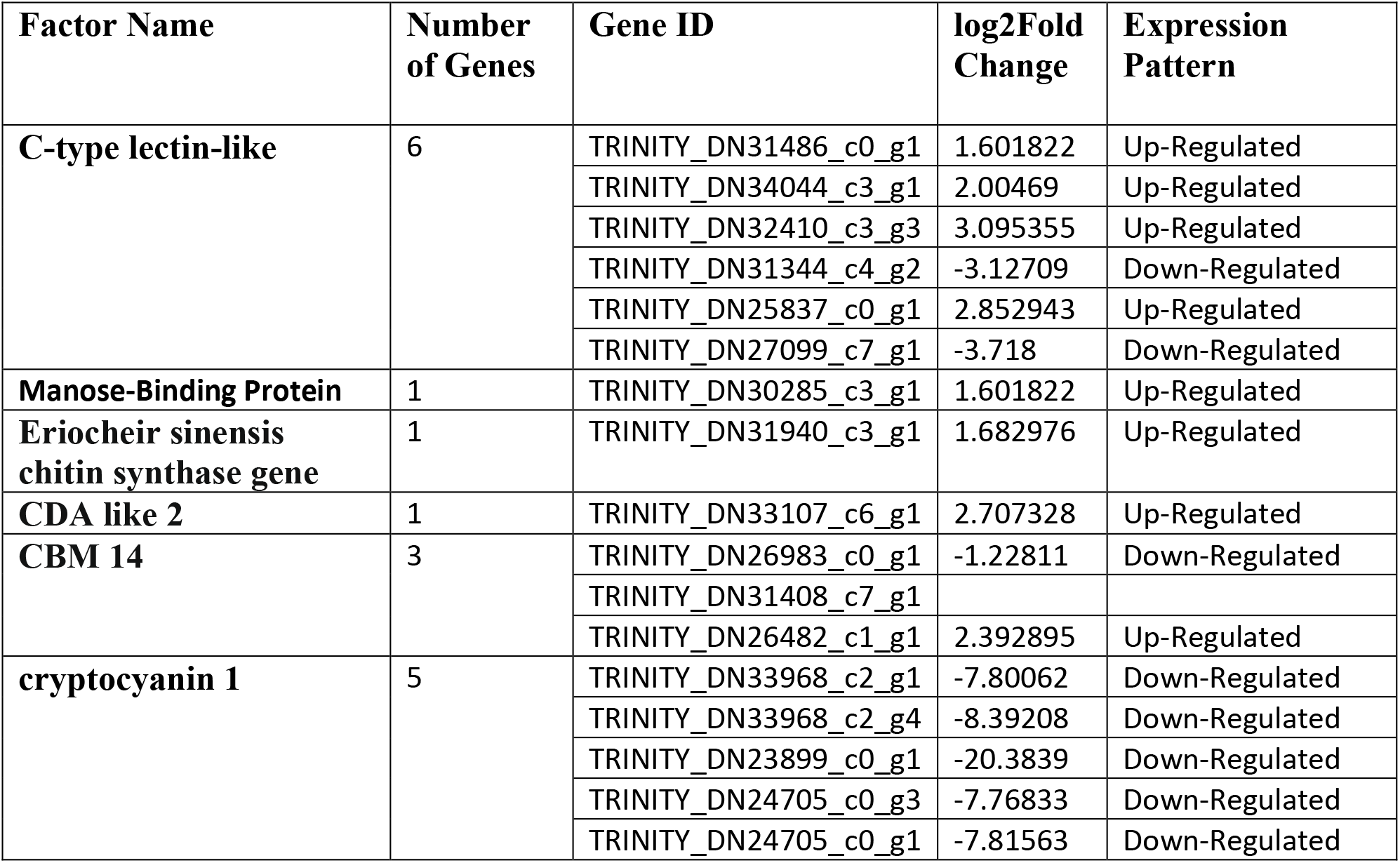
Exoskeleton Related Factors

### 3.7 Analysis of Transcriptome Data by qRT-PCR

To validate the RNA-seq findings, we chose eight genes that were significant enriched in KEGG analysis of the M vs NM molting stages, and performed RT-PCR. The selected genes include MGAM, mfsd8, ACOX1, ninaB, Nanp, DCXR, CYP2L1, dhdh, NIL, CYP2L1 and AF436001 (internal reference gene). We found that results that RT-PCR were similar to those of RNA-seq (**Fig.** S4).

## 4. Discussion

Molting in crayfish is one of the biological processes, worth paying attention to, this because it is an important process that promotes growth, development, and reproduction (Chang, 1995; Raviv, et al., 2008; Chang & Mykles, 2011; Dai, 2017). Moreover, loss of the exoskeleton renders the animal vulnerable to infections and predation which in turn has significant implication in the aquaculture industry. The value of understanding the molecular mechanisms underlying the molting process of crayfish as well as of other decapods is appreciated for a variety of purposes including disease control studies, growth enhancement, exoskeleton formation and predation studies, among others (Devaraj & Natarajan, 2006; Kuballa & Elizur, 2008; Gore,2017). Nevertheless, the molecular mechanisms underlying crayfish molting process still remains poorly understood. Therefore, the present study was conducted to provide more insights in this area.

The molting process is divided into four continuous but distinct stages (intermolt, pre-molt, molt, and post-molt) (Skinner, 1985; Keller, 1992). In this study, analysis of the transcripts from the two molting stages (NM and M), divided them into two groups (Fig. 2). Showing a higher correlation between samples from each molting stage, hence, a big difference occurs between the two stages of the crayfish sampled for this study (Fig. 2c). In PCA plot, the proximity of the samples among the two groups show the similarity of gene expression among them in that molting stage. This indicates the phase of progression of the molting process in that molting stage (M and NM). In NM stage, the samples (NM1, NM 2 and NM 3) were widely sparsed showing that they were at different phases of progression of the NM stage, this was expected because the non-molt stage range from intermolt to premolt stage. On the other hand, samples in the M stage (M1, M 2 and M 3) were close to each other, indicating that the phase of progression was similar among them. A comparison of the gene expression profiles of the two molting stages, showed that post-molt (M) (percent) had highly differentially expressed genes compared to the intermolt stage (NM) (%), (Fig. 2b & Fig. 3a). This is because post-molt (NM) is a critical stage that may have more developmental and immune response genes involved (Wassenberg, & Hill, 1984; Chang, 1995).

### Energy Demand During the Molting Process

Factors related to mitochondrial proteins, such as cytochrome and NADH dehydrogenase were identified (Tab. 4; Tab. S2 & S3). NADH dehydrogenase and cytochrome are two of the three energy-transducing enzymes in the mitochondrial electron transport chain (Young, 1972; Vasudevan, & S, 2007; Navarro & Boveris, 2007). NADH dehydrogenase show up-regulation in the M (post-moult) compared to non-molt stage. The expression profile of NADH dehydrogenase seem to reflect an increase in the energy requirements of the animal in M stage. This is because in the post-molt stage the animal is recovering from molting, hence, a lot of metabolic activities are taking place, requiring a substantial amount of energy (Wassenberg, & Hill, 1984; Chang, 1995). Interestingly several studies have reported lower levels of energy requirements during molting, followed by a sharp rise in the post-molt stage due recovery in metabolic activity and returning to normal in the intermolt and the rest of the other molting stages (Chang, 1995; Kuballa et al., 2011). Cytochrome, however, was down-regulated in the M stage when compared to the NM stage. In animals, Cytochrome and NADH dehydrogenase have been shown to be stimulated by thyroid hormones through the stimulation of mitochondrial activity (Enríquez, et al., 1999). However, in this study the thyroid hormone signaling pathway was down-regulated in M stage when compared to the NM stage. (ko04919; Thyroid hormone signaling pathway).

Furthermore, phosphagen kinases such as creatine kinase and arginine kinase were identified and found to be significantly upregulated in M stage compared to NM stage (Tab. 4). Phosphagen kinases serve in intracellular energy transport and as temporal ATP buffer (Eggleton, 1934; Suzuki, & Furukohri, 1994; Ellington, 2001). Phosphagen kinases is found in different parts of crustaceans, however, they are more abundant in muscles and gills (Ellington, 2001). Arginine kinase activity was found to vary significantly based on the tissue in *Carcinus maenas*, in which its abundance corresponded with energy requirements of the tissue (Kotlyar, et al., 1999). Therefore, given the fluctuation in energy demands during the molt cycle, we therefore assume that phosphagen kinases plays a key role as ATP buffer to in order meet these fluctuating energy demands.

### 4.2 Immuno-regulation Process during molting

Crustaceans have quite a lot of defense mechanisms that become activated depending on the stage of the molting process or the pathogen’s characteristics (Hoffmann, 1999; Iwanaga, & Lee, 2005; Vazquez, et al., 2009). The defense mechanisms of crustaceans totally depend on the innate immune system, activated when pathogen related molecular patterns are recognized by soluble or by cell surface proteins of the host, lectins, antimicrobial e.t.c, these in turn, activate humoral or cellular effector mechanisms to destroy invasive pathogens (Vazquez, et al., 2009). For this study, of particular importance was to learn the crayfish immune mechanism during the non-molting stage (inter-molt-pre-molt) and the post-molt period. In crustaceans, several researchers have found the molt and post-molt stages as periods when the animal is more susceptible to pathogenic infections and stress (Xu, et al., 2020). This is largely because, its new exoskeleton is not yet well developed, a lot of physiological processes are taking place and has a low exercise capacity. In the present study several immune related factors were identified and were found to be highly up-regulated (molt vs nonmolt camparisons) (Tab. 5; Tab. S2 & S3).

The GO term representing (GO:0045087), innate immune response was highly expressed in the M stage, representing an activated immunity to counter possible invasion during the stage. Based on the KEGG pathway analysis for DEGs between the M and NM, some signaling pathways related to innate immune system of invertebrate were identified. Lysosome was one of the most significantly enriched KEGGY pathways (Fig. 4a). Activities related Lysozymes (GO:0003796) a product of lysosome, were found to be highly expressed in the post molt stage. The Lysozymes are widely distributed immune effectors involved in numerous physiological processes, such as in digestive systems and immune (Van Herreweghe, 2012; Rowley, 2016; Chen, et al., 2018) exerting cytosolic activity on peptidoglycans of bacterial cell walls to initiate cell lysis. Moderate Lysozyme antimicrobial activity were revealed in mud crab (*S.paramamosain*) (Zhou et al., 2017). Toll-like receptor was another signaling pathway found to be significantly up-regulated in M stage compared to the NM. Toll-like receptors (TLRs) are a group of molecules that play an essential role in the recognition of Pathogen-related molecular patterns (PAMPs) and in the initiation of innate immune responses to infectious substances (Kawai, 2005; Marion, 2018; Sánchez-Paz & Muhlia-Almazán, 2020).

Caspase, interluking 1 beta converting enzyme was upregulated in N vs M, thus highly expressed in the post-molt satge. Interluking 1 beta enzyme is a cysteine protease that converts pro-IL-1β into active IL-1β. IL-1β is a pro inflammatory cytokine that mediates many of the physiological and behavioral responses to inflammation (Nakajima & Iwamoto; 2011; Gu et al., 2017). A study conducted recently showed that IL-1 family may play an important role in activating innate immune responses against pathogen infection in mud crab (Gu et al., 2017). Generally, NOD-like Receptors (NLRs) detect intracellular bacterial and viral infections and induce the inflammasome complex, which leads to caspase-1-dependent processing and secretion of members of the interleukin 1 (IL-1) family of cytokines, such as IL-1β and IL-16 (Franchi et al., 2009).

Serine proteases (SPs) are among the biggest enzyme families in the animal kingdom and play essential roles in immune responses (Shi, et al., 2008). In this study, Trypsin-like serine protease was up-regulated in the M. In crustaceans, trypsin-like serine proteases have been isolated from the hepatopancreas of the Chinese shrimp (*Fenneropenaeus chinensis*) and the red claw crayfish (*Cherax quadricarinatus*), were observed to involved in the innate immune defense against pathogens (Shi, et al., 2008; Fang, et al., 2013). Serine proteinase homologs were also obtained from mud crab (*S. paramamosain*) and it has been suggested that *Sp*-SPH protein could bind to a number of bacteria and play a key role in host defense against microbe invasion (Zhang et al., 2013).

Generally, just before molting, during molting, and after molting there is an increase in the metabolic activities in crustaceans, leading to additional burden on the antioxidant system, which in turn increases the risk of oxidative stress (Alcaraz & Sard’a 1981; Penkoff & Thurberg 1982; Cockcroft & Wooldridge 1985). Generally, oxygen consumption rises, reaching a peak shortly before ecdysis, and then declines rapidly after molting to increase again to basal levels in the intermolt stage (Filho et al., 2001). Several, antioxidant enzymes are secreted during molting, these enzymes help in compensating for the oxidative stress caused by metabolism or invading microorganism (immuneresponse) (Liu, et al., 2010). Here we found that redoxin enzyme family including peroxiredoxin, thioredoxin and glutaredoxin proteins were significantly up-regulated in N vs NM comparison. Thioredoxin, with a redox active disulfide bridge, is essential for sustaining the balance of reactive oxygen species, which has vital role on the immune system (Koháryová & Kollárová, 2015; Cunningham, et al., 2015). Thioredoxin has been characterized in mud crab and it is believed to be a potential biomarker gene for environmental stress evaluation in marine species (Hu, et al., 2014). The peroxiredoxins (Prxs) define a novel and evolutionarily conserved superfamily of peroxidases able to protect cells from oxidative damage by catalyzing the reduction of a wide range of cellular peroxides (Wood et al., 2003; Rhee et al., 2005). Six peroxiredoxins were recently characterized in mud crab (Tu et al., 2018). They were also characterized in crayfish during a bacteria challenge and other crustaceans (Wu et al., 2017). Superoxide dismutase was also significantly upregulated in the M stage compared to the NM period. Superoxide dismutase is an enzyme that helps break down potentially harmful oxygen molecules in cells (Rhee et al., 2005; Tu et al., 2018).

C-type lectins (carbohydrate binding proteins) were down-regulated in M stage compared to NM stage. C-type lectins are involved in immune function through the lectin-complement pathway, where mannose-binding lectin identifies infectious substances and in turn activates the PO system (Decker, et al., 2007; Zhang, et al., 2015).

### 4.3 Exoskeleton Calcification and Sclerotization Related Activities

Hardening of the new exoskeleton is a complex process which involves calcification (mineralization) and sclerotization (Richards, 1951; Neville, 1975). Two factors implicated in the control of calcification in diverse structural matrices of invertebrates, C-type lectin-like domain and mannose-binding protein were found to be significantly upregulated in the post-most (M) compared to Non-molt stage (NM) (Tab. 6 and Tab. S3). C-type lectin-like domain and mannose-binding proteins have been shown to be relevant to the molt cycle-related modification of cuticular glycoproteins (Matsushita & Fujita, 1996). Glycosylation of cuticular proteins is believed to be a controller of biomineralization of the exoskeleton of crustacean (Shafer, et al., 1995; Tweedie et al., 2004). It has been suggested that glycoproteins serve as premolt inhibitors of calcification in the exocuticle, and that deglycosylation post-molt may eliminate this barrier to mineralization (Tweedie et al., 2004). C-type lectin-like is implicated in the inhibition of calcification by glycosylation of cuticular proteins, since its expression coincided with the formation of the new cuticle in the pre-molt and must remain uncalcified until the post-molt (Kuballa & Elizur, 2008). The C-type lectin was found to be highly expressed in the pre-molt and dropped as the molt proceeded through to post-molt stage and eventually reached lower levels in the intermolt stage. Similarly, our results show higher expression of c-type lectin in M stage compared to NM. Since, we did not sample crayfish in premolt stage we cannot really determine the expression pattern of the C-type lectin across the moltage, nonetheless its expression in the M stage could be attributed to the inhibition of calcification of the membranous layer, arthrodial membranes, gills and the gut which remain uncalcified at all the times. The up-regulation of C-type lectin in the M stage can also be attributed to fact it plays a role in immune responses, as described earlier.

The expression of mannose binding protein, on the other hand has been attributed to the facilitation of calcification of the exoskeleton through deglycosylation of cuticular proteins (Akita et al., 1992). Kuballa & Elizur (2008), found that up-regulation of mannose binding protein was specific only to the post-molt period, at the time when cuticle calcification takes place, strongly suggesting its involvement in calcification process. In this study, expression of mannose binding protein was more expressed in the M stage than the NM, hence, we postulate that mannose binding protein calcification of the exoskeleton.

To further understand the activities related to the exoskeleton formation, the expression pattern of other factors such as Chitin synthase, CDA, CBM, and Cryptocyanin were evaluated (Tab. 6). Chitin synthase (CHS) is an essential membrane protein in arthropods, responsible for the synthesis of chitin from UDP-N-acetylglucosamine and its transfer to the extracellular domain (Merzendorfer, 2003). So far two chitin synthases are recognized in insects, (Arakane et al., 2004; Hogenkamp et al., 2005), of which one has been shown to be involved in cuticle construction in larvae and adult arthropod peritrophic membrane. In our results, chitinase, was upregulated in the M stage when compared to the NM stage indicating its potential involvement in the formation of the new exoskeleton. CDA is one of the exoskeleton related protein identified in this study, it was found to be significantly upregulated in the M stage. Generally, CDA converts chitin into chitosan, by digesting chitin acetyl groups. Hence, it has an important role in exoskeleton formation. Tom and Collegues (2014), annotated two CDAs of which one was prominently expressed. Furthermore, cryptocyanin, was involved in the formation of the new exoskeleton in crustaceans, was identified (Terwilliger, e al., 2005). However, it was down regulated in the M stage compared to the NM stage. We also found that chitin binding peritrophin-A domain (CBM 14) was up-regulated in the M stage compared to NM stage. CBM 14 is found in chitin binding proteins, chiefly as peritrophic matrix proteins of insects, these proteins function of the peritrophic membrane and are important in the structural formation in insects (Terra, 2001). Up-regulation of CBM 14 in the M stage, reveals its involvement in cuticle synthesis and hardening.

### 4.4 Hormone Regulation Process during Molting

Crayfish molting process is controlled by a complex interplay of hormones that work together or independently throughout the molting cycle (Pamuru et al., 2012). The signals that come by neuropeptide hormones activate a series of physiological activities associated to molting, including losing the extracellular cuticle, escaping from the confines of the cuticle relatively rapidly, taking up water, expanding the new flexible exoskeleton and then quickly hardening it for defense and locomotion (Skinner, 1985). In our results several neuropeptide hormones and hormone related genes were identified and analyzed (Tab. 7).

It is well recognized that CHH and MIH peptide families are key hormones controlling the molting process in crustaceans (Chung & Webster, 2003). MIH, secreted by a neuro-secretory center known as the X-organ/sinus gland complex (XO/SG) in crustacean eyestalks, belongs to the CHH family. Ablation of XO/SG has been shown to reduce MIH secretion, which in turn enable ecdysteroid synthesis, and enhanced molt frequency (Pamuru et al., 2012). MIH was up-regulated in the stage compared to NM stage (Tab. S3), indicating that it was highly expressed in the M stage compared to NM stage (intermolt to pre-molt). This was also observed in the study conducted by Qiao et al (2018) in *V. nipponense*, MIH expression increased rapidly to the highest level in the post-molt and remained lowly expressed in intermolt stage and reached it’s lowest levels in pre-molt stage. Similarly, Huang et al. (2014), in a study on the green mud crab *Scylla paramamosain reported similar findings*. This may be due to the fact that the animal has just undergone molting and may not need to undergo molting just as soon, hence, the hormone is secreted in high level. Moreover, it marks the beginning of the new molt cycle. In this study we observed that growth gonad inhibiting hormone (GIH) up-regulated (Tab), exhibiting a molting pattern similar to that of MIH, hence, we assumed that they may have common functions. Similarly, Qiao et al (2018) reported that GIH and MIH were closely related, and postulated that they have similar functions.

To further understand hormones downstream the molting process of Red swamp crayfish pathway and clarify possible roles in this process, we enumerated several other hormonal-related transcriptional factors of the two molting stages. According to the results, a trend of up-regulated transcript levels was observed in majority of the identified factors during M stage compared to NM stage (Tab. 7). Ecdysteroid regulated-like protein, ecdysteroid kinase and E75 nuclear receptor were reported previously to be associated with molting; and they have been found to related to hormonal control during molting (Nakagawa & Henrich, 2009). In this present study, ecdysteroid regulated-like protein, ecdysteroid kinase and E75 nuclear receptor were highly expressed in the postmolt in the postmolt (M), suggesting that they could be involved in hormonal regulation during this stage. The vitellogenin gene (VG) is believed to be is an ecdysteroid reactive gene; in *Cherax quadricarinatus*, synthesis of the vitellogenin gene has been revealed to be induced by 20E during the premolt (Souty-Grosset, 1997). In this study, we observed a down-regulated trend of vitellogenin gene in N vs NM. Neuroparsin, were first recognized as anti-gonadotropic factors which delay vitellogenesis in insects (Girardie, et al., 1987, 1989). The inhibitory effect of neuroparsin on vitellogenesis and oocyte maturation was confirmed by RNA interference in female *Schistocerca gregaria* (Badisco, et al., 2011). In contrast, neuroparsin were observed to play a positive role in the maturation of oocytes by igniting vitellogenin production in the hepatopancreas (Yang, et al., 2014). In this study, two genes encoding neuroparsin were identified, one was upregulated while the other was down-regulated.

## Conclusion

Tracing the expression patterns of genes during molting can help in explaining and deducing the molecular mechanisms involved in the molting process. The information presented here provides a basic understanding of the molecular mechanisms underlying the crayfish molting process, with respect to energy requirements, hormonal regulation, immune response, and skeletal related activities during post-molt stage and the intermolt stage. In the present study more genes and other factors were detected, however, we mainly focused on highly expressed factors and those described in other studies as relevant to energy requirements, hormonal regulation, immune response, and exoskeleton in arthropods with much focus on those found in crustaceans. The expression pattern of these genes in the two molting stages is evident that molting is a complex process encoded by a large number of gene families, involving many mechanisms and requiring strict control system.

## Acknowledgements

This work was supported by grants from the National Nonprofit Institute Research Grant of CATAS-TCGRI (2019JBFM01). We thank the students and staff of the Aquatic Genetic Laboratory, FFRC for their assistance in the study.

## Author Contributions

Shengyan Su, Yongkai Tang and Pao Xu conceived the study, Brian Pelekelo Munganga contributed to designing of the experiments. Brian Pelekelo Munganga, Fukuan Du, Juhua Yu, JianLin Li, Fan Yu, Meiyao Wang, Xinjin He, Xinyuan Li, Raouf Bouzoualegh performed red swamp crayfish data collection. All authors contributed to drafting the manuscripts.

---

**Figure S1.**
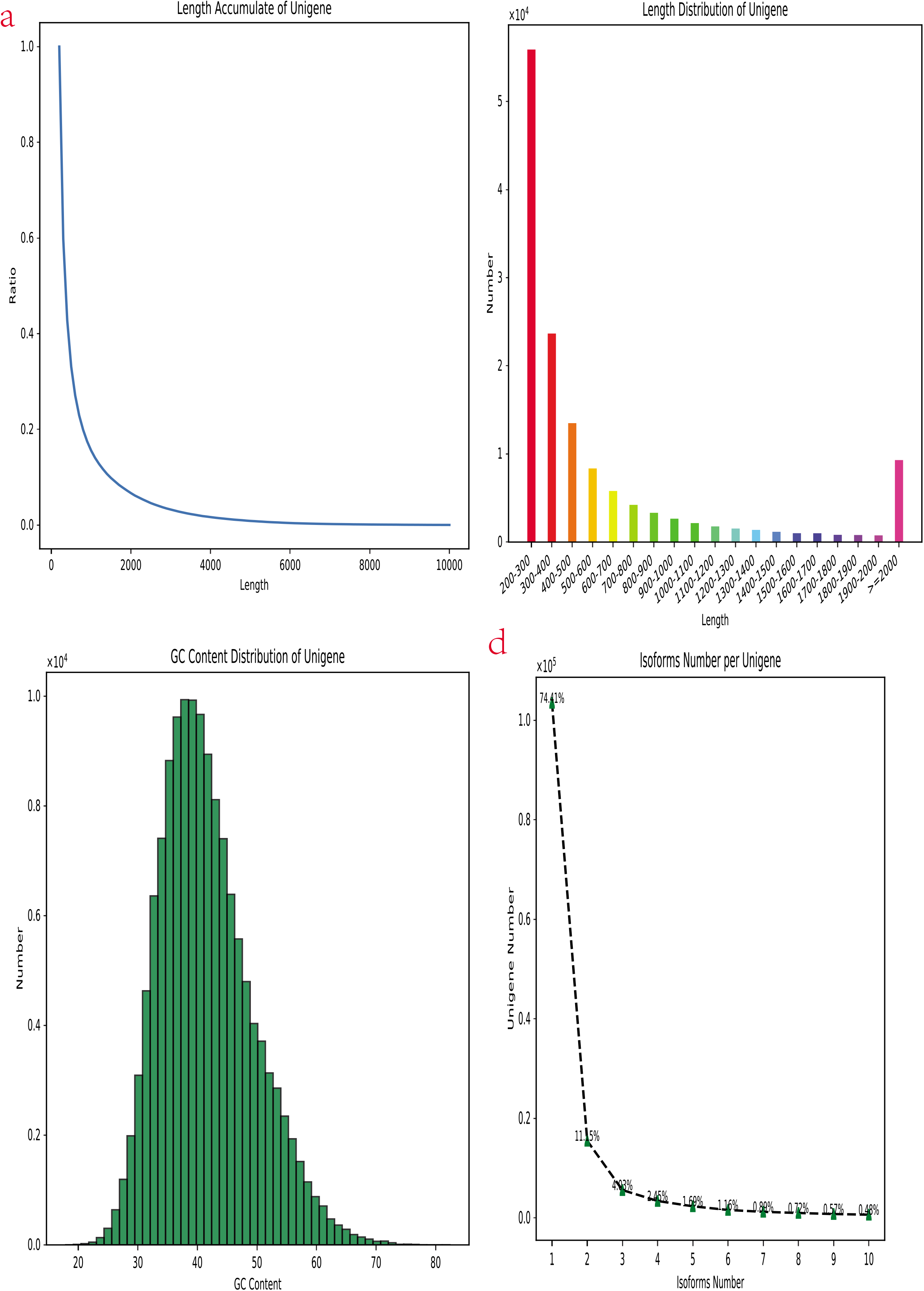
(a) Bar graph of unit size against SSR number per Mbp of M vs NM. (b) Statistics of single nucleotide polymorphisms (SNPs) and insertion/deletions (indels) in the transcriptome (N vs M).

**Figure S2.**
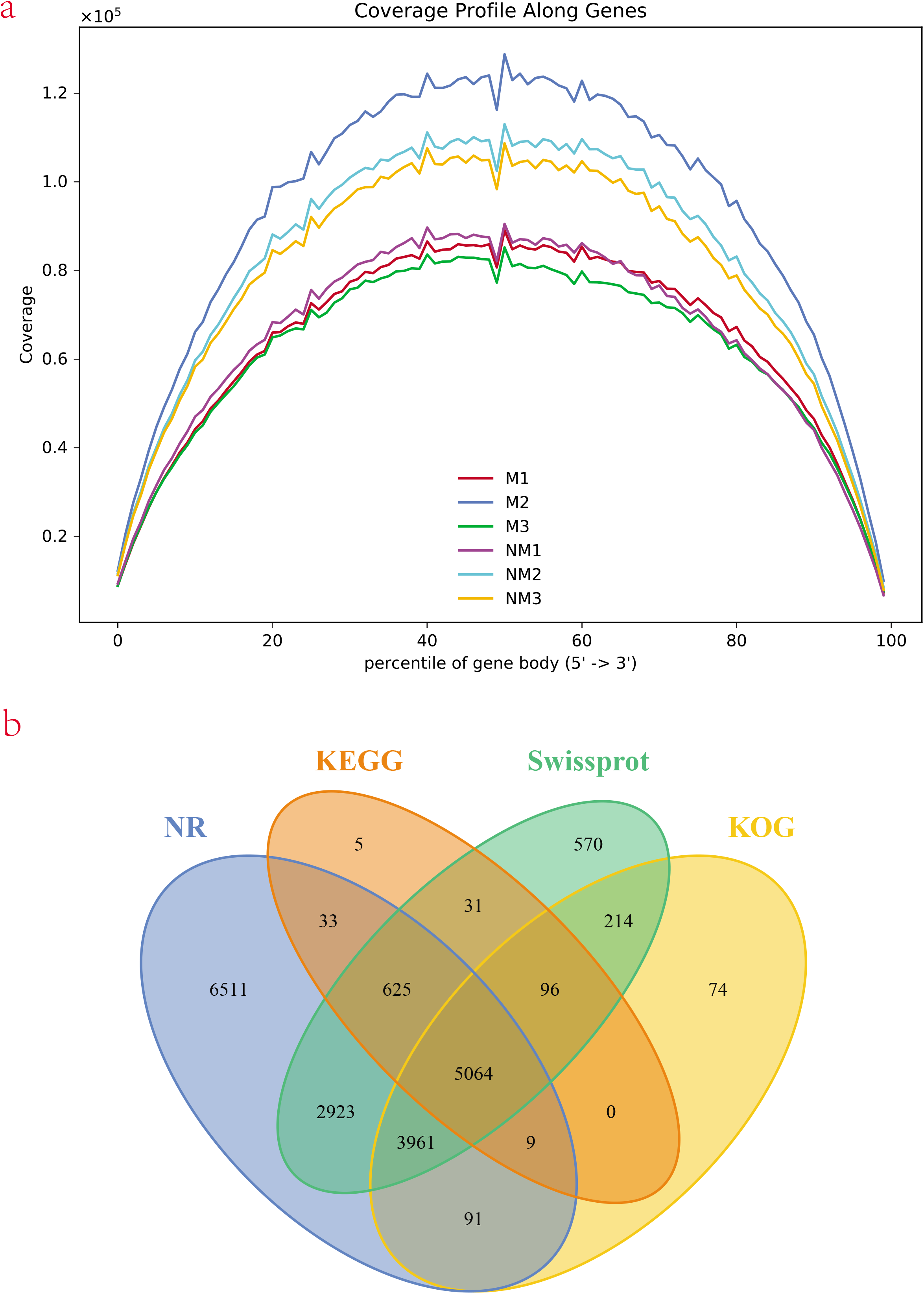
(a) Reads coverages of gene body percentile. (b)Venn diagram of annotation results against NCBI, NR, KEGG, Swissprot, and KOG. The number in each color block indicates the number of unigenes that is annotated by single or multiple databases.

**Figure S3.**
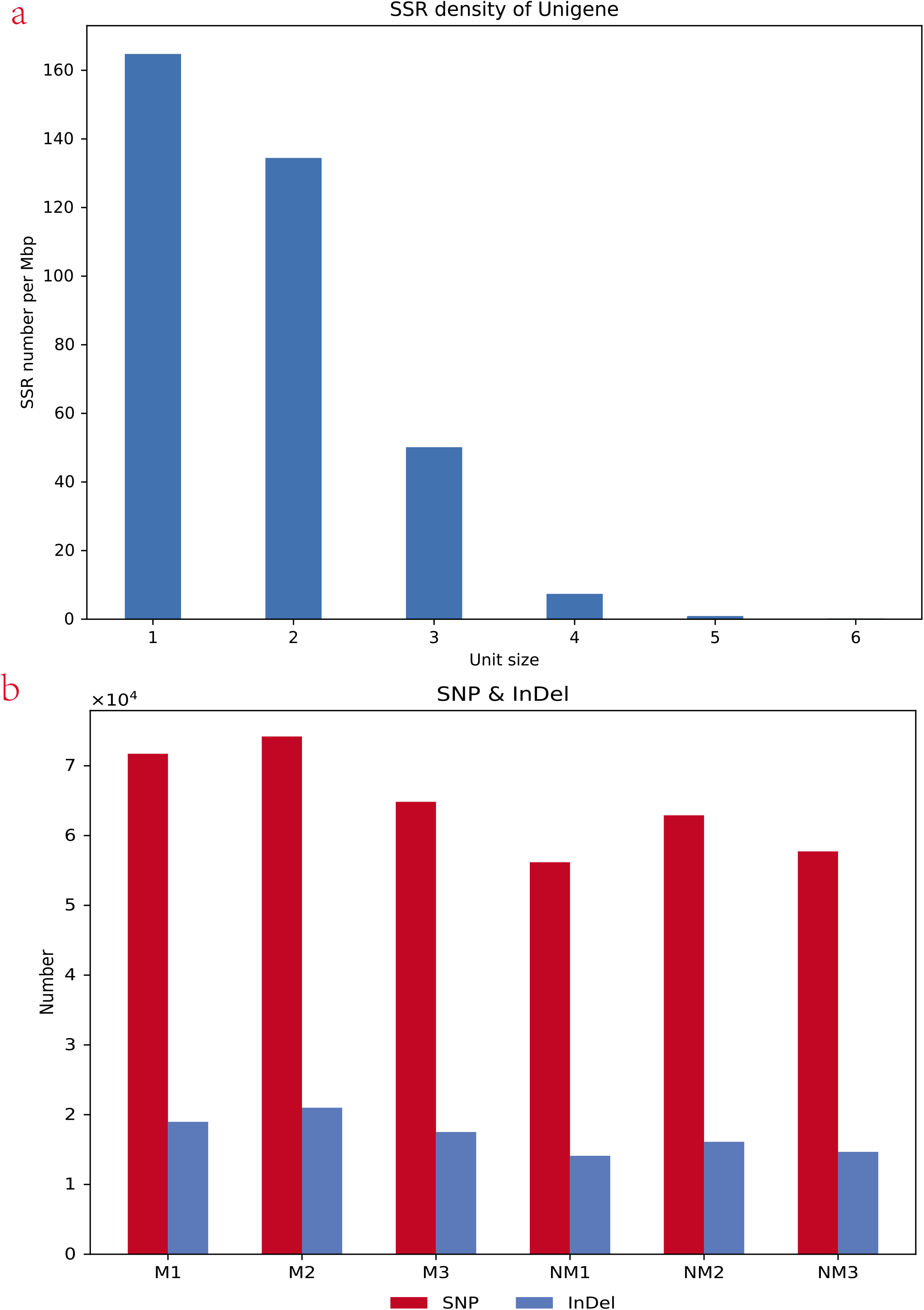
Length and GC-content of All-Unigene. (a) Bar chart depicting length distribution of All-Unigene identified in this study (length vs ratio of unigenes). (b) Bar chart depicting length distribution Bar chart depicting length distribution (length vs number unigenes). (c) GC content frequency distribution of All-Unigene of this study. (d) Isoform distribution of the unigenes.

|  | NM2 | NM3 | NM1 | M1 | M3 | M2 |
| --- | --- | --- | --- | --- | --- | --- |
| Total Reads | 52358598 | 50324500 | 43018686 | 46972334 | 46798082 | 68136198 |
| Total Bases | 7.61E+09 | 7.27E+09 | 6.23E+09 | 6.79E+09 | 6.81E+09 | 9.9E+09 |
| Average Read Length | 145.3 | 144.43 | 144.87 | 144.58 | 145.59 | 145.35 |
| Q10 Bases | 7.61E+09 | 7.27E+09 | 6.23E+09 | 6.79E+09 | 6.81E+09 | 9.9E+09 |
| Q10 Bases % | 100.00% | 100.00% | 100.00% | 100.00% | 100.00% | 100.00% |
| Q20 Bases | 7.49E+09 | 7.16E+09 | 6.14E+09 | 6.69E+09 | 6.75E+09 | 9.77E+09 |
| Q20 Bases % | 98.48% | 98.48% | 98.45% | 98.48% | 99.07% | 98.64% |
| Q30 Bases | 7.2E+09 | 6.88E+09 | 5.89E+09 | 6.43E+09 | 6.6E+09 | 9.41E+09 |
| Q30 Bases % | 94.63% | 94.64% | 94.54% | 94.61% | 96.88% | 95.03% |
| N Bases Count | 2331 | 2248 | 1972 | 2022 | 52894 | 5878 |
| N Bases Ratio | 0.00% | 0.00% | 0.00% | 0.00% | 0.00% | 0.00% |
| GC Bases Count | 3.72E+09 | 3.59E+09 | 2.97E+09 | 3.17E+09 | 3.19E+09 | 4.65E+09 |
| GC Bases Ratio | 48.86% | 49.35% | 47.62% | 46.67% | 46.83% | 46.91% |

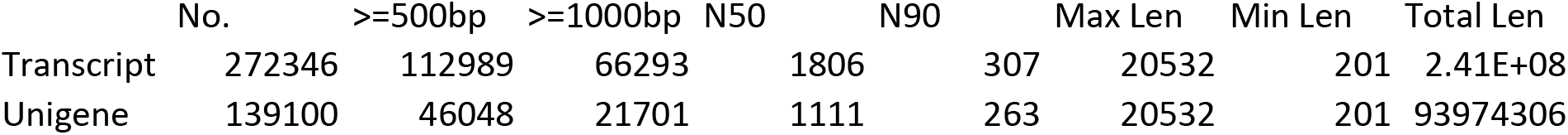

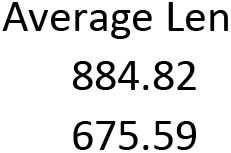

|  | M1 | M2 | M3 | NM1 | NM2 | NM3 |
| --- | --- | --- | --- | --- | --- | --- |
| Total reads | 45257896( | 65217100( | 44811426( | 41766832( | 51699876( | 49737936(100.00%) |
| Total mapped | 42935008( | 62234555( | 42017582( | 39907772( | 49559754( | 47640782(95.78%) |
| Multiple mapped | 34454998( | 50113055( | 33566048( | 31981708( | 40050309( | 38751145(77.91%) |
| Uniquely mapped | 8480010(1. | 12121500( | 8451534(1. | 7926064(1. | 9509445(1. | 8889637(17.87%) |
| Read-1 mapped | 4248326(9 | 6070241(9 | 4236539(9 | 3973519(9 | 4763168(9 | 4453249(8.95%) |
| Read-2 mapped | 4231684(9 | 6051259(9 | 4214995(9 | 3952545(9 | 4746277(9 | 4436388(8.92%) |
| Reads mapped | 4263747(9 | 6081211(9 | 4242478(9 | 3984923(9 | 4773963(9 | 4463769(8.97%) |
| Reads mapped | 4216263(9 | 6040289(9 | 4209056(9 | 3941141(9 | 4735482(9 | 4425868(8.90%) |
| Non-splice | 8480010(1. | 12121500( | 8451534(1. | 7926064(1. | 9509445(1. | 8889637(17.87%) |
| Splice read | 0(0.00%) | 0(0.00%) | 0(0.00%) | 0(0.00%) | 0(0.00%) | 0(0.00%) |
| Reads mapped | 7462780(1 | 10876092( | 7414518(1 | 6795300(1 | 8244322(1 | 7705816(15.49%) |

